## Supplementary Information for "Pollinators in food webs: Mutualistic interactions increase diversity, stability, and function in multiplex networks"

Supplementary Information includes:

S1: Plant-pollinator network construction.

S2: Uncertainty and sensitivity analyses.

S3: Feedback and other controls.

Table S1: Qualitative effects of variation in parameter values on persistence, abundance, and productivity.

Table S2: Parameters and quantitative results of uncertainty analyses.

Table S3: Parameters and quantitative results of sensitivity analyses.

Table S4: Summary of symbols, definitions, and values used in the main text.

Figure S1: Effects of mutualism on plant persistence in complex ecological networks.

Figure S2: Robustness of results to simulation length.

Figure S3: Connectance of network treatments.

Figure S4: Properties of empirical and simulated plant-pollinator networks.

Figure S5: Persistence along a gradient of increasing floral rewards productivity.

Figure S6: Illustration of controls.

Figure S7: Diversity, stability, and function of feedback controls.

Figure S8: Changes in community composition changes in feedback controls.

Figure S9: Example timeseries comparing multiples and feedback control simulations.

Figure S10: Diversity, stability, and function in rewards partitioning controls.

Figure S11: Changes in community composition changes in rewards partitioning controls.

**S1: Plant-pollinator network construction.**

To synthesize network structure in our multiplex model, we attempted to create empirically-realistic food webs and plant-pollinator networks that could be interconnected. This required us to choose a common definition of nodes for these networks, as nodes in empirical plant-pollinator networks often represent taxonomic species while nodes in empirical food webs are often represented as "trophic species," in which all taxonomic species with similar sets of predators and prey are grouped into one node (Cohen & Briand 1984, Martinez 1991, Williams & Martinez 2000). Because trophic-grouping can reduce methodological bias in empirical food web data and additionally facilitate comparison to structural models (Dunne 2006), we followed a similar approach to aggregate species in empirical plant-pollinator networks ($N$ = 49 networks listed in Table S1 of Valdovinos *et al.* 2016). Specifically, we grouped species with exactly the same set of mutualistic partners into one node. This tended to group super-generalist plants and specialist pollinators into single nodes, increasing connectance ($C_p$), reducing species diversity ($S_p$), and thus reducing the average ratio of animal to plant species ($A/P$) of empirical networks from 2.5 to 2 (Fig. S4a). The 95% confidence intervals around the diversity-connectance relationship in the grouped empirical networks were defined by the following equations: $C_{p\_min} = 1.05811 S_p^{-0.6572}$, $C_{p\_max} = 2.1737 S_p^{-0.4779}$, where $C_{p\_min}$ bounds the minimum connectance and $C_{p\_max}$ bounds the maximum connectance across a range of diversities.

We generated plant-pollinator networks within this range of empirical properties using the stochastic mutualistic network algorithm from Thébault and Fontaine (2010, main text Fig. 2b). The algorithm takes as input the number of pollinators ($A$), the number of plants with pollinators ($P$), the network connectance ($C_p = L_p / AP$, where $L_p$ is the number of pollination links), and additional shape parameters *pcomp* and *pnest*. We used $P$ = 3, 4, …, to 19 plants with pollinators, corresponding to 15 to 95% of the original 20 plants in the niche-model food webs (main text Figs. 2a, 2c). For simplicity, we fixed the ratio of animal-to-plant species to the value derived from the aggregated empirical networks ($A/P = 2$), yielding plant-pollinator network diversities of $S_p = A + P = 9, 12, \ldots,$ to 57. For each $S_p$, we subdivided the range of connectances between $C_{p\_min}$ and $C_{p\_max}$ into 10 intervals, giving 11 values for $C_p$, inclusive of endpoints. Parameters *pcomp* and *pnest* are probabilities that determine rules by which plants and pollinators are linked together (see Thébault & Fontaine 2010, Supplementary Online Methods). Increasing *pcomp* towards 1 leads to more *modular* networks, i.e. networks where groups of species interact more among themselves than with species from other groups. Increasing *pnest* towards 1 leads to more *nested* networks, i.e. networks wherein specialist species tend to interact with partners that are a subset of the partners of more generalized species. Empirical mutualistic networks tend to be highly nested (see *Network properties*, below), but this property varies substantially between similar empirical networks and is strongly related to their diversity and connectance. We ran Thébault and Fontaine's (2010) algorithm 10 times for each of the 11 $C_p$'s corresponding to each $S_p$, with *pnest* varied from 0, 0.2, …, to 1 and *pcomp* fixed to 0 for a total of 10*11*17*5 = 9,350 networks. Only those networks in which each species had at least one link were retained. In particular, the confidence intervals for low $S_p$ networks include values for $C_p$ in which connectance is too low for all species to have at least one partner; these networks were excluded. From the remaining networks, we chose networks at each diversity level that most evenly sampled the empirical range of nestedness and connectance (Fig. S4a-b). This resulted in approximately 14 plant-pollinator networks at each of the 17 initial diversity levels ($S_P = 9, 12, \ldots, 57$), for a total of 238 networks.

*Network properties*

Nestedness is a frequently observed, but controversial property of empirical mutualistic networks that has been hypothesized to endow them with both stability and function (Valdovinos *et al.* 2016). However, its effect appears to be dependent on both model assumptions (Bascompte *et al.* 2003, Okuyama & Holland 2008, Allesina & Tang 2012, James *et al.* 2012, Rohr *et al.* 2014, Valdovinos *et al.* 2016, Pascual-García & Bastolla 2017) and its formal definition including its potential correlation to degree heterogeneity (Saavedra & Stouffer 2013). We therefore quantified both nestedness and degree heterogeneity of our simulated plant-pollinator networks (Fig. S4b).

Specifically, we quantified nestedness as overlap and decreasing fill (NODF), standardized to the mean NODF of 100 iterations of the "CE" null model in the ANINHADO software package (Guimarães & Guimarães 2006). This null model creates randomized matrices with the probability of a link between $i, j$ set to the mean fraction of links realized in the $i$th row and $j$th column of our simulated interaction matrix. Standardized nestedness ranges from -1.0 to 1.1 in our simulated networks.

We defined "degree heterogeneity" as the standard deviation of the degree distribution of all species (plants and pollinators), so that a more positive value indicates a more heterogeneous network. Degree heterogeneity ranges from 0.67 to 5.21 in our simulated networks and is significantly correlated with nestedness (R = 0.829, N = 238, $P < 0.0001$), as expected.

Modularity is also a potentially stabilizing property of ecological networks though it tends to be inversely related to nestedness and is more often observed in herbivory networks than in mutualistic networks (Thébault & Fontaine 2010). We quantified bipartite modularity in our plant-pollinator networks using the Newman's leading eigenvector method ($Q$, Newman 2006) standardized by the mean of 100 iterations of the "Fixed" null model in the BiMat software package (Flores *et al.* 2015). This null model swaps rows and columns of our simulated interaction matrix to create randomized matrices with the exact same row and column sums. Standardized modularity ranges from -0.34 to 0.64 in our simulated networks and is negatively correlated with nestedness ($R = 0.451$, $N = 238$, $P < 0.0001$).

*References*

Allesina, S. & Tang, S. (2012). Stability criteria for complex ecosystems. *Nature*, 483, 205-208.

Bascompte, J., Jordano, P., Melian, C.J. & Olesen, J.M. (2003). The nested assembly of plant-animal mutualistic networks. *Proc. Natl Acad. Sci. USA*, 100, 9383-9387.

Cohen, J. E. & Briand, F. (1984). Trophic links of community food webs. *Proc. Natl. Acad. Sci. USA*, 81, 4105-4109.

Dunne, J.A. (2006). The network structure of food webs, in *Ecological Networks Linking Structure to Dynamics in Food Webs*, pp. 27-86. Oxford U. Press, Oxford.

Flores, C.O., Poisot, T., Valverde, S., & Weitz, J.S. (2015) BiMat: A MATLAB package to facilitate the analysis of bipartite networks. *Methods in Ecol. Evol.*, 7. 127-132.

Guimarães, P. R. & Guimarães P. (2006). Improving the analyses of nestedness for large sets of matrices. *Environ. Modell. Software* **21**, 1512-1513.

James, A., Pitchford, J.W. & Plank, M.J. (2012). Disentangling nestedness from models of ecological complexity. *Nature*, 487, 227-230.

Martinez, N. D. (1991). Artifacts or attributes? Effects of resolution on the Little Rock Lake food web. *Ecol. Monogr.*, 61, 367-392.

Okuyama, T. & Holland, J.N. (2008). Network structural properties mediate the stability of mutualistic communities. *Ecol. Lett.*, 11, 208-216.

Newman, M.E.J. (2006). Modularity and community structure in networks. *PNAS*, 103 (23), 8577-8582.

Pascual-García, A. & Bastolla, U. (2017) Mutualism supports biodiversity when the direct competition is weak. *Nat. Comm.*, 8:14326, 1-13.

Rohr, R.P., Saavedra, S. & Bascompte, J. (2014). On the structural stability of mutualistic systems. *Science*, 25, 416-425.

Saavedra, S. & Stouffer, D.B. (2013). "Disentangling nestedness" disentangled. *Nature*, 500, E1-E2.

Thébault, E. & Fontaine, C. (2010). Stability of ecological communities and the architecture of mutualistic and trophic networks. *Science*, 329, 853-856.

Valdovinos, F.S., Brosi, B.J., Briggs, H.M., Moisset de Espanés, P., Flores, J.D., Ramos-Jiliberto, R. & Martinez, N.D. (2016). Niche partitioning due to adaptive foraging reverses effects of nestedness and connectance on pollination network stability. *Ecol. Lett.*, 19, 1277-1286.

Williams, R.J. & Martinez, N.D. (2000). Simple rules yield complex food webs. *Nature*, 404, 180-183.

**S2: Uncertainty & sensitivity analyses.**

We explored the behavior of our multiplex model and assessed its robustness by performing uncertainty and sensitivity analyses on its free parameters (Table S4). For these analyses, we simulated the $N$ = 24,276 networks from each of the Rewards Only (RO) Food Web (FW), RO multiplex, and Rewards Plus (RP) multiplex treatments for 2000 timesteps, which was approximately dynamical steady-state (Figs. 3, S2).

For the uncertainty analyses (Table S2), we varied only one parameter varied at a time (Table S2) with all other parameters fixed to the main text values recorded in Table S4, but with two exceptions: the half-saturation density fixed to the same value for all consumers ($B_0$ = 60), and the rewards productivity rate fixed to High ($\beta$ = 1), unless otherwise noted. The qualitative effect of variation in each parameter is summarized in Table S1 and quantitative results are given in Table S2. A particularly important result, the effect of rewards productivity rate ($\beta$) on species persistence, is shown in Fig. S5. For the sensitivity analyses (Table S3), we varied all parameters simultaneously using a Latin Hypercube Sampling (LHS) design. Following Thébault & Fontaine (2010), we varied the parameters between ¼ to 4 times the value used in the main text (Table S4), but with $\beta$ = 0.6 and $B_0$ = 60 for all consumers.

We additionally explored the effect of basic network structural properties on persistence in our six main text treatments. Specifically, we applied a Classification and Regression Tree (CART) analysis with five-folded cross-validation to predict species persistence at the end of simulations given the following metrics of initial network structure (Berlow *et al.* 2009): network treatment (RO or RP), rewards productivity level ($\beta$ = 0 [FW], 0.2, 1), initial diversity ($S$), connectance of the network ($C$), connectance of the niche-model food web ($C_f$), connectance of the plant-pollinator network ($C_p$), nestedness of the plant-pollinator network (NODFst and degree heterogeneity), and modularity of the plant-pollinator network (*Q* and *Qst*). All inputs are continuous variables except for network treatment and rewards productivity level, which are categorical. See S1 and Table S4 for definitions of structural metrics.

*Results*

As reported in other work (Williams & Martinez 2004, Martinez *et al.* 2006), the form of the functional response for feeding (main text Eqn. 3) as controlled by the Hill coefficient (*h*) and the half-saturation density ($B_{0ij}$) can modify the diversity-persistence relationship in complex food webs (Tables S1-S3). Otherwise, our results are most strongly mediated by food availability for pollinators (Table S1), which is directly affected by rewards productivity ($\beta$, Fig. S5) and rewards self-limitation (*s*) rates and indirectly affected by the cost of rewards production to vegetative total growth ($\kappa$) and community-wide carrying capacity of plant vegetation (*K*). In general, increasing rewards availability *directly* (increasing $\beta$ and/or decreasing *s*) increases species persistence (e.g., Fig. S5), pollinator biomass, and biomass of omnivores and carnivores so that they exceed that of FWs and furthermore allows for a positive diversity-persistence relationship. *Indirectly* increasing rewards availability (decreasing $\kappa$ and/or increasing *K*) increases persistence, but changes the shape of how biomass accumulates in ecosystems with increasing initial diversity so that the diversity-persistence relationship shifts to negative; this may be an example of the paradox of enrichment.

In general, multiplex networks above a certain threshold of food availability for pollinators displayed increased persistence and biomass compared to FW treatments, indicating that this result is robust to variation in parameter values, though the qualitative relationship between initial diversity (*S*) and species persistence may vary. The effects of food availability for pollinators are most obvious in the Rewards Only (RO) multiplex treatment, where increasing food availability increases pollinator abundance and subsequently the abundance and persistence of omnivores and carnivores. On the other hand, in the Rewards Plus (RP) multiplex treatment, increasing food availability increases pollinator abundance, which increases the abundance and persistence of omnivores and carnivores but additionally leads pollinators to exert increased direct competition and predation pressure on lower trophic-level consumers (herbivores and omnivores), reducing species persistence and abundance in the ecosystem overall. However, the RP multiplex treatment has increased persistence and abundance at low pollinator food availability than analogous RO networks. See Fig. S5 for an example of this pattern driven by rewards productivity ($\beta$). An interesting possibility suggested by these results is that the increased trophic links in RP networks may dampen the effects of mutualistic feedbacks, whether they are stabilizing or destabilizing. This hypothesis is also coherent with the results of our feedback control (S3).

Regardless of the choice of parameters, multiplex networks tend to exhibit higher total biomass and productivity than the RO FW treatment, and increasing rewards availability tends to correspond to an increasingly positive relationship between initial diversity ($S$) and species persistence.

Furthermore, CART analyses on all $N$ = 145,656 simulations in our six main text treatments underscore the importance of rewards productivity ($\beta$) for species persistence in our networks. Rewards productivity accounts for 73% of the explained variance (five-folded $R^2$ = 0.48, $N$ = 145,656), while network connectance ($C$) and initial diversity ($S$) account for ~9% each, and other network metrics including nestedness and modularity account for < 2% each. Subdividing our networks by initial diversity ($S$) yields similar results. For example, when $S$ = 64, rewards productivity accounts for 73% while $C_f$ and $C$ each account for ~11% of the explained variance in persistence (five-folded $R^2 = 0.37$, $N = 8568$). When $S = 80$, rewards productivity accounts for 85% and $C$ accounts for 10% of the explained variance in persistence (five-folded $R^2 = 0.61$, $N = 9180$). In both cases, metrics of nestedness and modularity explain < 4% of explained variance in observed species persistence.

*References*

Berlow, E. L. *et al.* (2009) Simple prediction of interaction strengths in complex food webs. *Proc. Natl Acad. Sci. USA,* 106, 187–191.

Martinez, N. D., Williams, R. J., Dunne, J. A. & Pascual, M. (2006) Diversity, Complexity, and Persistence in Large Model Ecosystems, in *Ecological Networks Linking Structure to Dynamics in Food Webs*, pp. 163–184. Oxford U. Press, Oxford. doi:10.1111/j.1469-7998.1982.tb03499.x

Thébault, E. & Fontaine, C. (2010). Stability of ecological communities and the architecture of mutualistic and trophic networks. *Science*, 329, 853-856.

Williams, R. J. & Martinez, N. D. (2004). Stabilization of chaotic and non-permanent food-web dynamics. *Eur. Phys. J. B,* 38, 297–303.

**S3: Feedback and other controls.**

Multiplex treatments had higher average diversity, persistence, biomass, productivity, consumption, and stability (lower temporal variability) than their counterpart FW treatments (main text Fig. 4), with the exception of the Low Rewards Only treatments which displayed lower average diversity, persistence, productivity, and species-level stability. Which mutualistic mechanisms cause the diversity, stability, and function in our multiplex model to differ from traditional food webs? In particular, what is the role of floral rewards, a resource available to consumers in the multiplex but not the food web (FW) treatments?

At steady-state of the multiplex simulations, plants' production of rewards interacts with their vegetative production and their pollinators' consumption. These interactions emerge from the dynamic feedbacks between plants and pollinators whereby plants produce rewards, which pollinators consume while providing reproductive services, which increases vegetative growth rate, which affects vegetative biomass, which affects rewards productivity, etc. (see Fig. S6a). To understand the role of rewards biomass in the multiplex model, we created two dynamic control treatments by forcing non-mutualistic systems to produce rewards at rates of a steady-state mutualistic system, but without the mutualistic feedbacks. The feedback control is applied at initialization to understand if mutualistic feedbacks, isolated from rewards availability, shape final ecosystem diversity, stability, and function during transient dynamics. The rewards partitioning control is applied at steady-state to understand if the mutualistic dynamics and/or the resource partitioning of rewards and vegetation (i.e. dividing plants with pollinators into two nodes) hold the ecosystem at steady-state. This allows us to test whether the additional biomass produced by plants with pollinators solely leads to the observed stability and function of our multiplex networks or whether mechanisms of mutualism are required for these effects. These two feedback controls also represent two ways that traditional food web models could be re-parameterized to accommodate rewards biomass and productivity without the added complexity of the mutualistic dynamics in the multiplex model. If the control simulations yield different results than the multiplex simulations, we can conclude that mutualistic dynamics are important flows to quantify in order to make accurate predictions in empirical systems.

*Feedback control*

Our feedback control (Fig. S6b) transforms species of plant with pollinators into two *independent* biomass pools: a vegetation pool and a rewards pool. Productivity of the pools is fixed to the mean rates of the analogous nodes in a multiplex network. This severs mutualistic feedbacks operating during the simulations so that vegetation and rewards production are no longer interrelated, pollinators provide no reproductive services to plants when consuming their rewards, and plants' growth rates are independent of pollinators' behavior. However, the rates or productivity that emerged from mutualism in the non-control multiplex networks and the resource partitioning between vegetation and rewards is preserved. See main text Methods: Feedback control for details.

We compared the results of these simulations with those of the original multiplex simulations by measuring absolute differences in persistence and total biomass at timestep 5000, where the *effect of feedback = multiplex – control*. To assess differences in these community metrics due to guilds, we calculated absolute differences in the fraction of persisting species composed by each guild:

$$\frac{multipex\ final\ guild\ diversity}{multiplex\ final\ diversity} - \frac{control\ final\ guild\ diversity}{control\ final\ diversity} \qquad \text{(S1)}$$

and the fraction of biomass composed by each guild:

$$\frac{multipex\ guild\ biomass}{multiplex\ total\ biomass} - \frac{control\ guild\ biomass}{control\ total\ biomass}. \qquad \text{(S2)}$$

When these differences are positive, feedbacks in multiplex simulations have a positive effect, i.e. they increase persistence or biomass of the community or guild of that in the controls. When these differences are negative, feedbacks decrease persistence or biomass. We additionally computed all metrics of stability and function applied to the main text simulations.

*Rewards partitioning control*

To disentangle the influences of *rewards partitioning* and *mutualistic feedbacks* on our results, we created a rewards partitioning control (Fig. S6c) that transform plants with pollinators into a *single* biomass pool that includes both vegetation and rewards production but with a constant amount of biomass (equal to the final rewards biomass of the analogous species in a multiplex network) excluded from plant competitive dynamics, mimicking the steady-state multiplex dynamics. Specifically, we recorded average net rewards production ($\overline{\beta_i B_i - s_i R_i}$), vegetative biomass ($\bar{B_i}$), rewards biomass ($\bar{R_i}$), and realized growth rate ($\hat{r_i} = \overline{r_i P(R_i)}$) for each plant with pollinator *i* during the last 1000 timesteps of our multiplex simulations. Then at timestep 4000, we set the biomass of each plant with pollinators *i* to $\widehat{B_i} = B_i + R_i$ and their dynamics were modified to:

$$\frac{d\widehat{B_i}}{dt} = \left(1 - \frac{1}{K}\sum_{j \in plants}(\widehat{B_j} - \bar{R_j})\right)\hat{r_i}\widehat{B_i} - \sum_{j \in consumers} C_{ji}(\widehat{B_i})/e_{ji} + (1 - \kappa_i)(\overline{\beta_i B_i - s_i R_i}) \quad (5\text{-}1)$$

Eqn. 5-1 severs the dynamic feedback of mutualistic benefit due to provisioning of reproductive services to plants ($P(R_i)$ in Eqn. 5, Methods: Network dynamics) while preserving the total biomass of each plant (vegetation plus rewards), the realized growth rate of the plant ($\hat{r_i}$), the total rewards production rate ($(1 - \kappa_i)(\overline{\beta_i B_i - s_i R_i})$), and the exclusion of rewards from interspecific competitive interactions between plants $\left(\frac{1}{K}\sum_{j \in plants}(\widehat{B_j} - \bar{R_j})\right)$. As in the feedback control, it eliminates mutualistic feedbacks but additionally removes partitioning between vegetation and rewards, an important component from the perspective of consumers. Pollinators' consumption was necessarily switched from rewards to vegetative biomass so that, as in the FW treatments, pollinators' dynamics followed Eqn. 1 instead of Eqns. 1-1 or 1-2. All other species dynamics followed their original equations (Methods: Network dynamics). We then ran these simulations for an additional 1000 timesteps beginning with the biomass distribution at timestep 4000. For this analysis, we were most interested in whether the system was perturbed from steady-state by this change in dynamics even when productivities at the base of the food web were held constant.

*Results*

Overall, ecosystem diversity, persistence, biomass, and productivity in our feedback controls equilibrate to similar values as in the multiplex simulations (Fig. S7-8). The only overall difference was increased vegetative biomass of plants with pollinators and decreased biomass of plants without pollinators in the feedback controls (Fig. S8d). Though plants with pollinators in multiplex treatments can potentially achieve higher growth rates than plants without pollinators in the presence of sufficient reproductive services, plants with pollinators in feedback controls increase in biomass because they are not subjected to dynamic effects of mutualism such as variable vegetative growth rate and costly rewards production. All controls except High RP additionally displayed decreased omnivore persistence and biomass, an effect that was most prominent in the Low rewards controls (Fig. S8b, S8d). We also observed decreased herbivore and increased pollinator persistence in the RO controls and decreased added-omnivore/pollinator persistence in the Low RP control (Fig. S8b). These changes appear related to transient oscillations in species' biomass. The period and amplitude of pollinators' and plants' oscillations in the controls are less synchronized and decrease (e.g. Fig. S9) presumably due to steadier rewards biomass in the absence of its dynamic coupling to vegetation. However, omnivores, added-omnivores/pollinators in RP controls, and herbivores have increased oscillations with increased synchrony. We suggest that transient oscillations between mutualists may stabilize multiplex networks by increasing compensatory dynamics in consumers' resources which is especially important for species that integrate over many of the resources within an ecosystem such as omnivores. These guild-level differences were notable in the RO and Low RP treatments but tiny in the High RP treatment. This pattern suggests that the combination of sufficient rewards productivity and increased trophic connectedness of mutualists dampen mutualistic feedbacks. This is also suggested by the smaller ranges in outcomes between High and Low RP treatments compared to the larger range in High and Low RO treatments (Fig. 4, also see Fig. S5).

The pattern of similar ecosystem-level results in the multiplex and feedback control simulations but differences between them in guild persistence and abundance are coherent with the results of our rewards partitioning control. In this control, on average, species persistence (Fig. S11a) decreased by ~5-6% in RO and ~6-7% in RP treatments, while total community biomass (Fig. S11c) decreased by ~10% in High rewards treatments and ~6% in Low RP treatments but was unchanged in Low reward RO treatments. The magnitude of these reductions became larger as initial diversity increased, with the minor exception of total biomass in Low RO treatments (Fig. S10). Compared to the original multiplex simulations, total biomass in Low RO controls increased with initial increases of mutualism and diversity but then decreased precipitously with further increases, leading to no substantial change overall. Though these changes in community biomass and persistence are small overall, high variability and unbalanced production and consumption flows (Fig. S10b) indicate that turning "off" mutualism in the controls pushed the systems away from steady-state, demonstrating the importance of mutualistic feedbacks and/or rewards partitioning. More dramatically, the biomass and persistence of species and guilds changed in controls (Fig. S11b, d). Without partitioning of rewards from vegetation, former pollinators and herbivores could access to up to twice the biomass previously available to them. In response, pollinators decreased in abundance whereas herbivores increased in abundance but decreased in persistence. These changes combined with changes in consumers of species feeding on plants greatly change which species were more and less abundant in the controls while changing total community biomass relatively little. Thus, the dynamic feedbacks of mutualistic interactions, rather than solely increasing productivity at the base of the food web, increase persistence and biomass and substantially alter species abundance distributions in multiplex simulations. Additionally, these differences are exacerbated at higher levels of mutualism and diversity.

Our results suggest that the added productivity of mutualistic rewards primarily drive our observed patterns of ecosystem stability and function in the multiplex treatments (see S2 for analyses that further corroborate the role of rewards). However, our feedback control results also suggest that the *dynamics* of mutualistic feedbacks alter community composition in the ecosystem by increasing biomass and persistence of consumers, particularly omnivores, and decreasing biomass of plants with pollinators. Our rewards partitioning control results further suggests that resource partitioning between rewards and vegetation in plants with pollinators alters community composition by increasing biomass and persistence of consumers like herbivores that would otherwise face direct competition from pollinators.

In summary, diversity, stability, and function in multiplex ecological networks of food webs and pollination interactions result primarily from the added resource (floral rewards) it allows the system to create, but the dynamics of mutualism not only change the steady-state abundance distribution of species in the ecosystem through transient dynamics but also hold the system at that steady-state. If mutualistic dynamics were “turned off,” the system would be perturbed to a new steady-state abundance distribution with decreased persistence and abundance of herbivores and omnivores and increased abundance of plants. Our controls also illustrate that even if traditional food web models (like the dynamic model used for the Food Web (FW) treatments) were reparametrized to include rewards nodes with fixed productivity rates resulting from current mutualistic dynamics, such models would predict both transient dynamics and steady-state biomass distributions different from our multiplex model, which explicitly incorporates pollination dynamics.

**Table S1: Qualitative effects of variation in parameter values on persistence, abundance, and productivity.**

| Symbol | Main Text Value [Min., Max.] | Definition | Effect on Persistence | Observations & Hypothesized Mechanism |
|---|---|---|---|---|
| $\beta_i$ | Low = 0.2, High = 1; None corresponds to FWs [0.1, 1.8] | Production rate of plant with pollinator *i*'s floral rewards | ↑ | ↑ persistence & biomass of pollinators and their predators by ↑ food availability for pollinators; shifts multiplex networks to + diversity-persistence relationship |
| $s_i$ | 0.4 [0.1, 1.6] | Self-limitation rate of rewards production for plant with pollinator species *i* | ↓ | ↓ food availability for pollinators by slowing the recovery rate of depleted rewards, ↓ persistence & biomass of pollinators and their predators by ↑ competition among pollinators for resources |
| $\kappa_i$ | 0.1 [0.01, 1] | Cost of producing floral rewards biomass for plant with pollinator species *i* in terms of vegetative biomass | ~↓ | Stronger coupling of rewards production to vegetative growth rate; nonmonotonic effect on rewards but generally ↓ biomass of pollinators & plants with pollinators' vegetation, & ↓ pollinator persistence by ↓ their food availability |
| $B_{0ij}$ when *j* is species biomass | 60 [20, 100] | Half-saturation density; density of *j* at which *i* consumes at half its maximum feeding rate on *j* | ↓ | Slows consumption by *i* on rare resources, ↓ biomass & persistence of consumers at low diversity; shifts networks to + diversity-persistence relationship |
| $B_{0ij}$ when *j* is rewards | 30 [20, 100] | Same as above | ↓ | Slows consumption by pollinator *i* on rare rewards, ↑ rewards biomass, ↓ biomass & diversity of pollinators, especially at low diversity |
| $h$ | 1.5 [1, 2] | Hill coefficient, determines the shape of $F_{ij}$ | ↑ | Decelerates consumption on rare resources (Martinez *et al.* 2006); nonmonotonic effect on biomass, shifts networks to + diversity-persistence relationship |
| $K$ | 480 [120, 1320] | Plant community-wide carrying capacity | ↑ | ↑ basal food availability (vegetation); shifts networks to + then – diversity-persistence relationship |

Qualitative effects of each parameter as recorded from uncertainty analyses, where the focal parameter was increased from a Minimum to Maximum value (in brackets under the Main Text Value) while all other parameters were fixed. Patterns in persistence, biomass, production, and consumption when the parameter is increased are recorded as observations; hypothesized mechanisms leading to these patterns are also provided. If increasing the parameter modifies the qualitative relationship between initial diversity (*S*) and persistence, it is also recorded. Pollination parameters ($\beta$, $s$, $\kappa$, $B_{0ij}$ when *j* is rewards) are only relevant for multiplex networks. Remaining ($h$, $K$, and $B_{0ij}$ when *j* is not rewards) hold for all treatments.

**Table S4: Summary of symbols, definitions, and values used in the main-text.**

| Symbol | Definition | Main-Text Value | References |
|---|---|---|---|
| **Network architecture.** | | | |
| $S_f$ | Niche-model food web diversity (num. species) | 50 with exactly 20 plants and 5 herbivores | Fig. 1a |
| $L_f$ | Num. directed feeding links | $244 < L_f < 256$ | |
| $C_f$ | Niche-model food web directed connectance | $L_f / S_f^2 = 0.1$ | Martinez 1991 |
| $P$ | Num. plants with pollinator species | 3, 4, …, 19 | S1 |
| $A$ | Num. animal-pollinator species | $2P = 6$, 8, …, 38 | |
| $S_p$ | Plant-pollinator network diversity | $P + A = 9$, 12, …, 57 | Fig. 1b |
| $L_p$ | Num. directed pollination links | Range for each $S_p$ set by empirical networks | S1 |
| $C_p$ | Plant-pollinator network directed connectance | $L_p / PA$ = range for each $S_p$ set by empirical networks | |
| $S$ | Diversity of network treatments | $S_f + 2/3(S_p) = 56$, 58, …, 88 | Fig. 1c |
| $C$ | Directed connectance of network treatments | Range for each $S$ set by plant-pollinator network and inherited niche model links | Fig. 1d-e |
| **Network dynamics and parameterization.** | | | |
| $B_i$ | Biomass of species $i$ | Evaluated numerically from the system of differential equations | Eqns. 1, 4, 5 |
| $R_i$ | Floral rewards biomass of plant with pollinator $i$ | Same as $B_i$ | Eqn. 6 |
| $K$ | Plant community-wide carrying capacity | 480 | |
| $C_{ij}(B_j)$ or $C_{ij}(R_j)$ | Consumption rate of species $i$ eating species $j$ or $j$'s floral rewards, respectively | | Eqn. 2 |
| $F_{ij}$ | Functional response for $i$ eating $j$, describing the realized fraction of $j$'s biomass that is consumed as a function of $i$'s preference for $j$ and $j$'s prevalence | | Eqn. 3 |
| $\omega_{ij}$ | Preference of $i$ for eating $j$ | 1/($i$'s diet size), a.k.a. "the weak generalist model" | Williams 2008 |
| $h$ | Hill coefficient, determines the shape of $F_{ij}$ | 1.5 | Real 1977, Martinez *et al.* 2006 |
| $B_{0ij}$ | Half-saturation density; density of $j$ at which $i$ consumes at half its maximum feeding rate on $j$ | 60 when $j$ is species biomass, 30 when $j$ is rewards biomass | Boit *et al.* 2012 |
| $P(R_i)$ | Function describing plant with pollinator $i$'s accrual of *reproductive services* due to pollination | Of the form: *reproductive services* / (0.05 + *reproductive services*), where the shape parameter set to 0.05 is also called the *benefit coefficient* | Eqn. 7 |
| $\beta_i$ | Production rate of plant with pollinator $i$'s floral rewards | Rewards productivity treatments: Low = 0.2, High = 1 | |
| $s_i$ | Self-limitation rate of rewards production for plant with pollinator $i$ | 0.4 | |

| | | | |
|---|---|---|---|
| $\kappa_i$ | Cost of producing floral rewards biomass for plant with pollinator *i* in terms of its vegetative biomass production | 0.1 | |
| **Allometric parameterization.** | | | |
| $swTL_i$ | Short-weighted trophic level of species *i* | *i* is a plant: 1, herbivore: 2, omnivore or carnivore: >2 | Williams & Martinez 2004 |
| $x_i$ | Mass-specific metabolic rate of species *i* | *i* is a plant: 0, *i* is a consumer: $0.314m_i^{-0.25}$ | Brose *et al. 2006,* Martinez *et al.* 2012 |
| $m_i$ | Body mass of species *i* | $Z^{swTLi-1}$ with *Z* sampled from lognormal distribution with mean = 10, std. dev. = 100 | Martinez *et al.* 2012 |
| $r_i$ | Max. mass-specific growth rate of plant species *i* | Plants w/o pollinators: 0.8, plants w/ pollinators: 1 | Martinez *et al.* 2012 |
| $y_{ij}$ | Max. mass-specific consumption rate of *i* eating *j* | 10 | Martinez *et al.* 2012 |
| $e_{ij}$ | Assimilation efficiency of *i* eating *j* | *j* is a plant w/o pollinators or vegetative biomass: 0.66, *j* is an animal: 0.85, *j* is floral rewards: 1 | Martinez *et al.* 2012 |
| **Simulation settings.** | | | |
| *Initial Biomass* | Biomass of $B_i$ or $R_i$ at the beginning of the simulation | 10 | |
| *Simulation Length* | Num. timesteps in a simulation | 5000 | |
| *Solver* | Method used to numerically evaluate (“simulate”) differential equations specifying species’ dynamics | Multiplex: ode15s, Food Web (FW): ode45 used with $RelTol = 1^{-7}$, $AbsTol = 1^{-9}$ | MATLAB 2018b |
| *Extinction Threshold* | Biomass under which a species is considered extinct and its biomass is set to 0 for the remainder of the simulation | $10^{-6}$ | |

**Figure S1: Effects of increasing mutualism on plant persistence in complex ecological networks.**

Guild persistence of plants with (light blue) and without pollinators (dark blue), following the formatting of main text Fig. 5. Persistence is that fraction of initial species that persist (i.e. avoid extinction) to the end of simulations. Plants without pollinators always persist. Average persistence of plants with pollinators decreases with increasing diversity and mutualism, but overall extinctions are rare.

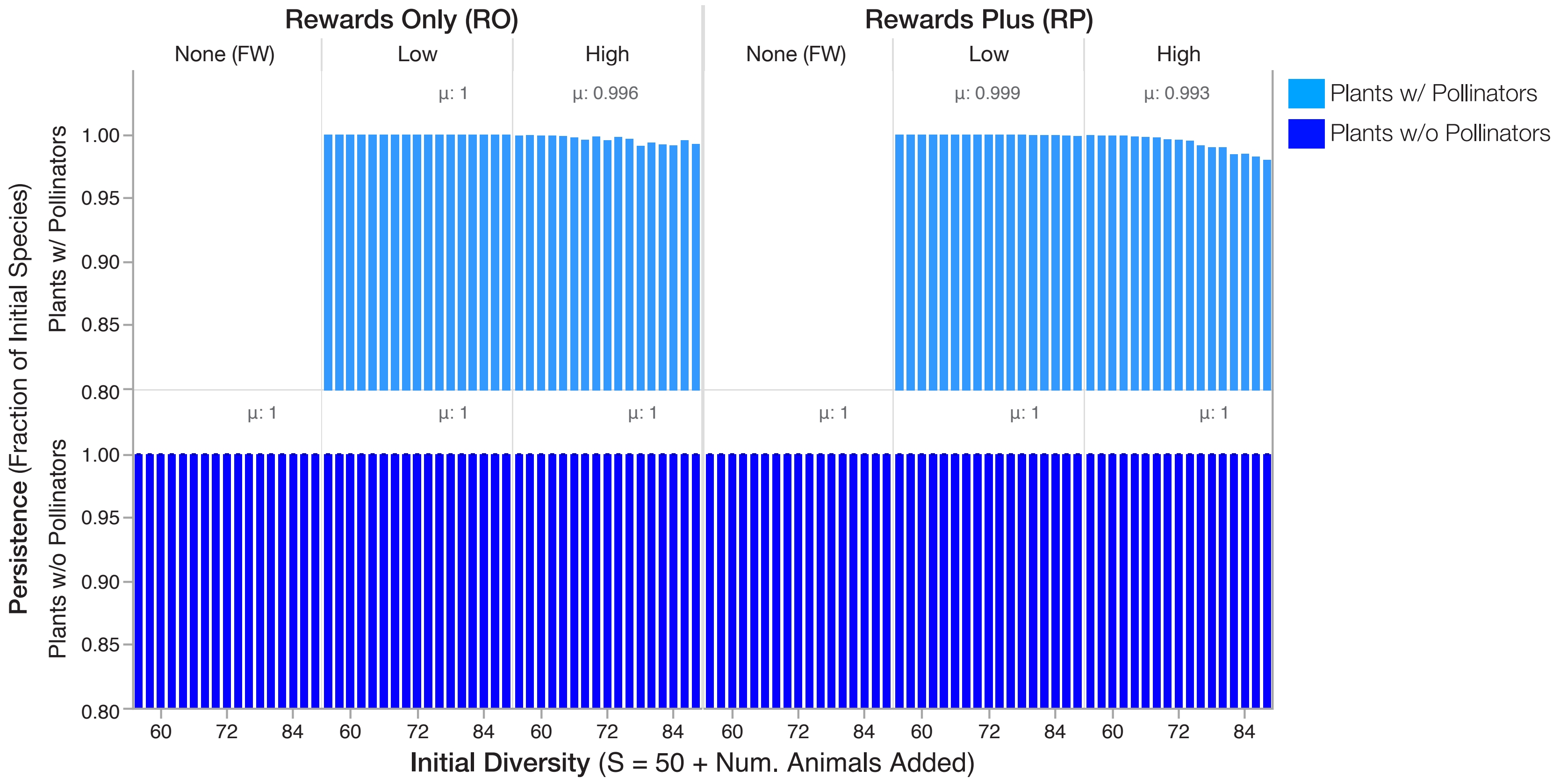

**Figure S2: Robustness of results to simulation length.**

Results of a subset of our simulations evaluated at different simulation lengths (x-axis). The simulations are of 90 randomly sampled $S = 72$ networks subjected to the six network treatments. Formatting of (a) and (b) follows main text Figs. 5 and 6, respectively. Hollow circles and diamonds indicate average values corresponding to the μ reported for species and guild CVs in Fig. 6e-f. Black vertical lines indicate results at 5000 timesteps, the length of the main text simulations.

**(a)**

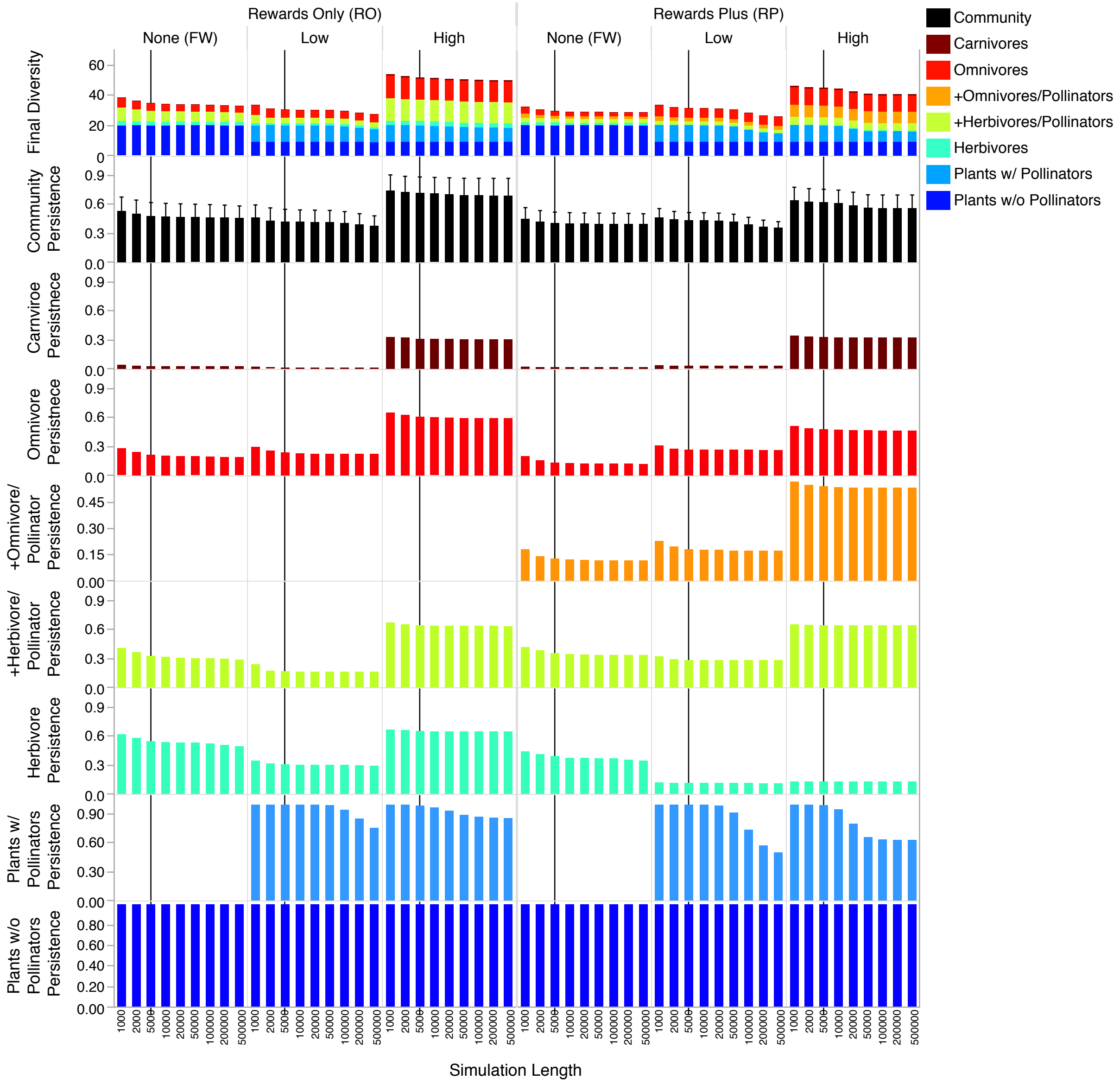

**(b)**

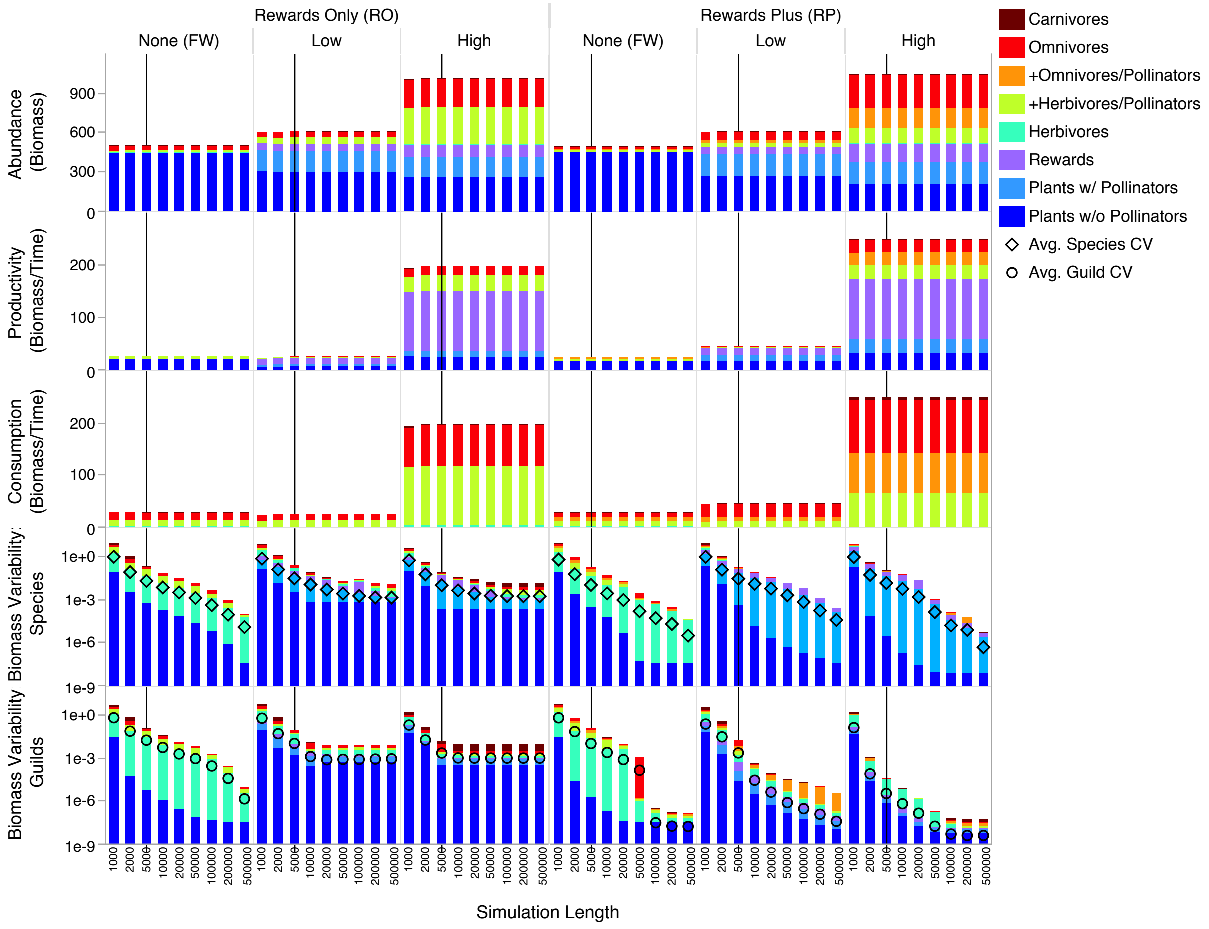
Rewards Only (RO)
Rewards Plus (RP)
None (FW)
Low
High
Abundance (Biomass)
Productivity (Biomass/Time)
Consumption (Biomass/Time)
Biomass Variability: Species
Biomass Variability: Guilds
Simulation Length
Carnivores
Omnivores
+Omnivores/Pollinators
+Herbivores/Pollinators
Herbivores
Rewards
Plants w/ Pollinators
Plants w/o Pollinators
Avg. Species CV
Avg. Guild CV

**Figure S3: Connectance of network treatments.**

The relationship of connectance (*C*) with species diversity (*S*) in our network treatments as described by linear regression (blue lines, given by the equation and $R^2$) and mean *C* across *S*. Connectance of the network treatments is defined as $C = L/S^2$, where *L* is the total number of links (both mutualistic and feeding) and *S* is the number of species in the network. Rewards Only (RO) treatments have identical connectance because every mutualistic link between pollinators and plants in the multiplex treatment is transformed into a feeding link by herbivores on plants in the Food Web (FW) treatment. Because the connectance of our simulated plant-pollinator networks decreases with increasing diversity (Fig. S5b) and pollinators have no other resources, *C* decreases with increasing *S* in the Rewards Only treatments. In the Rewards Plus (RP) multiplex treatment, pollinators can be both a pollinator and an herbivore of a given plant. These two links are transformed into a single herbivory link in the RP Food Web (FW) treatment, leading to slightly reduced connectance in the FWs (~ 0.10) compared to the multiplex networks (~0.11). RP pollinators were allowed to be herbivorous or omnivorous (main text Fig. 2d) such that *C* remains approximately constant with increasing *S* in RP FWs. *N* = 24,276 network in each treatment.

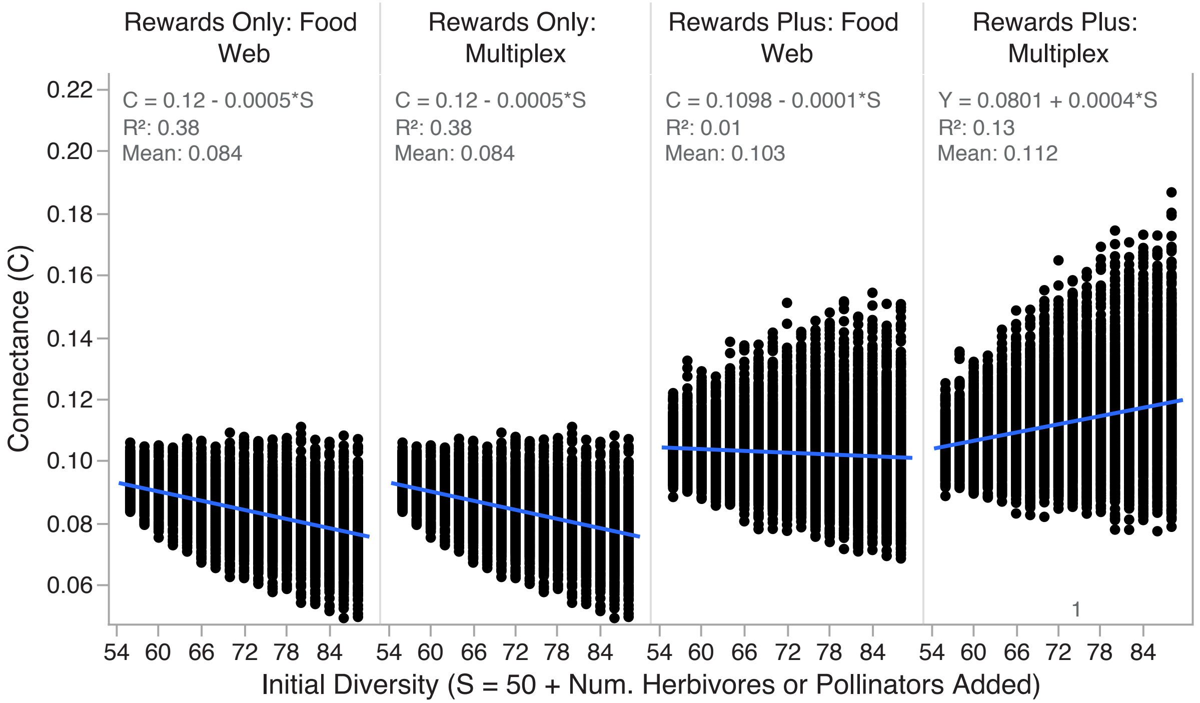

**Figure S4: Properties of empirical and simulated plant-pollinator networks.**

(a) The relationship of connectance ($C_p$) with species diversity ($S_p$) in $N = 49$ empirical plant-pollinator networks before (black, open circles) and after (blue, filled circles) trophic grouping and the 95% confidence intervals around each (lines). Simulated plant-pollinator networks (squares) have properties bounded by the grouped empirical networks. Note the log-log axes. (b) Properties of the 238 simulated plant-pollinator networks (squares). Connectance and diversity are highly correlated with nestedness and degree heterogeneity (colors).

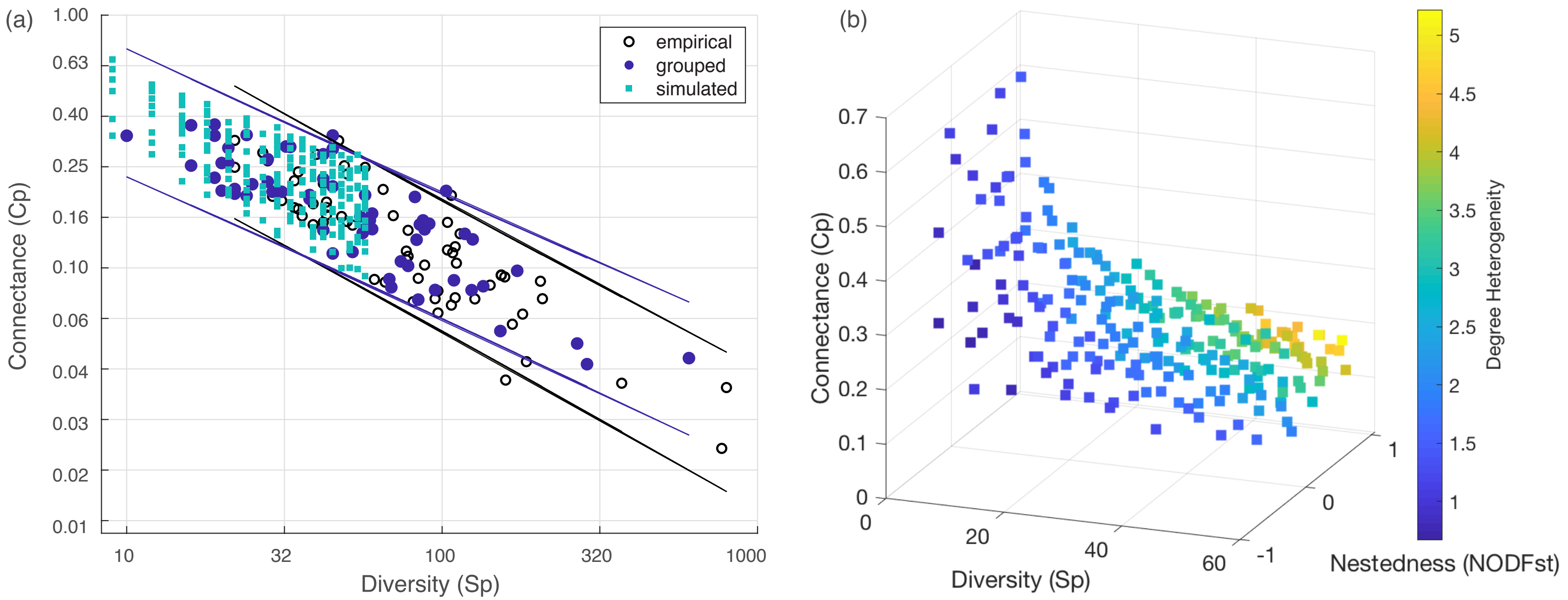

**Figure S5: Persistence along a gradient of increasing floral rewards productivity.**

Mean (bars) and standard deviation (error bars) of species persistence at varying rewards productivity for the $N$ = 24,276 networks each in the Rewards Only (RO) and Rewards Plus (RP) treatments. All others parameters were fixed (S1). The fraction of species that survive to the end of the simulations (persistence) is bounded between 0 and 1. At very low rewards productivity ($\beta$), persistence is ~0.4, corresponding to nearly all animals going extinct. Persistence increases with increasing $\beta$, but tends to level off so that there is a smaller difference in average persistence between $\beta$ = 1 and 1.4 than between $\beta$ = 0.8 and 1 (notice the non-linear x-axis). Persistence in RO networks (left) is more affected by $\beta$ than in RP networks (right).

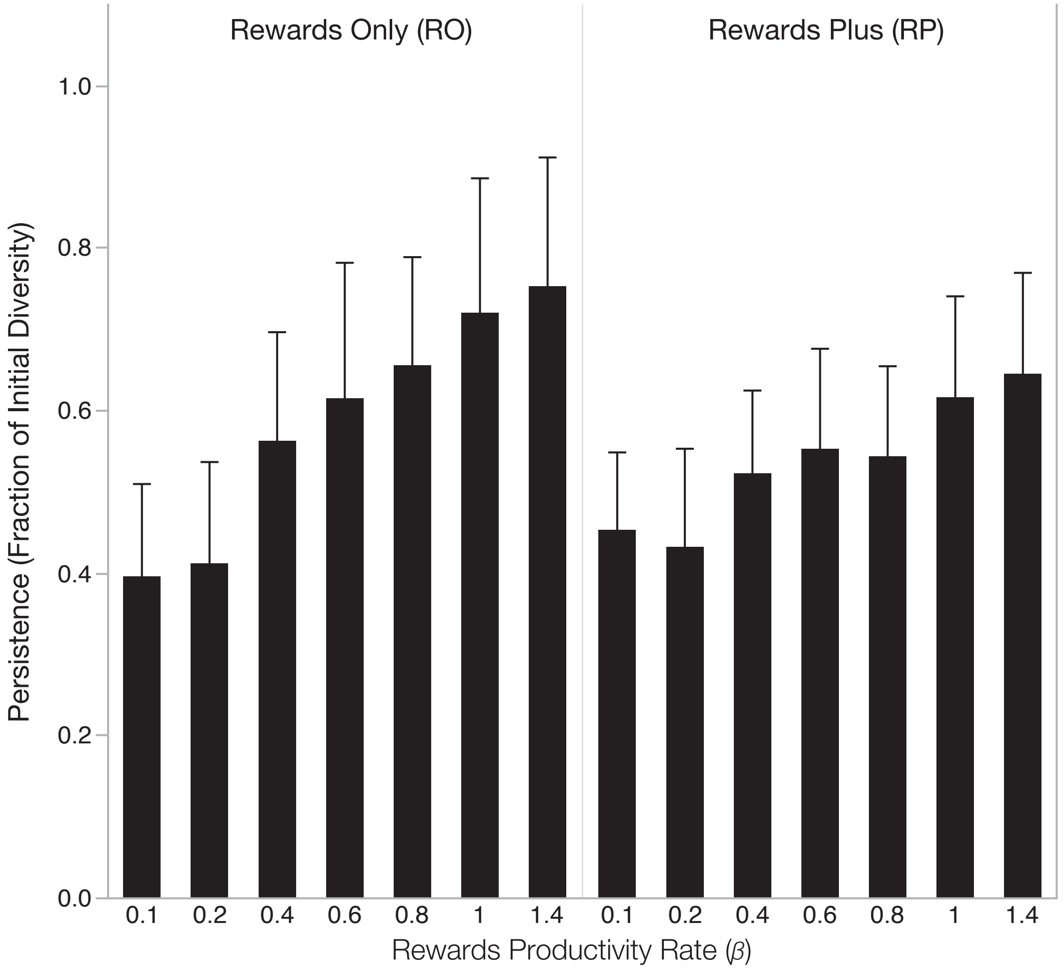

**Figure S6: Illustration of controls.**

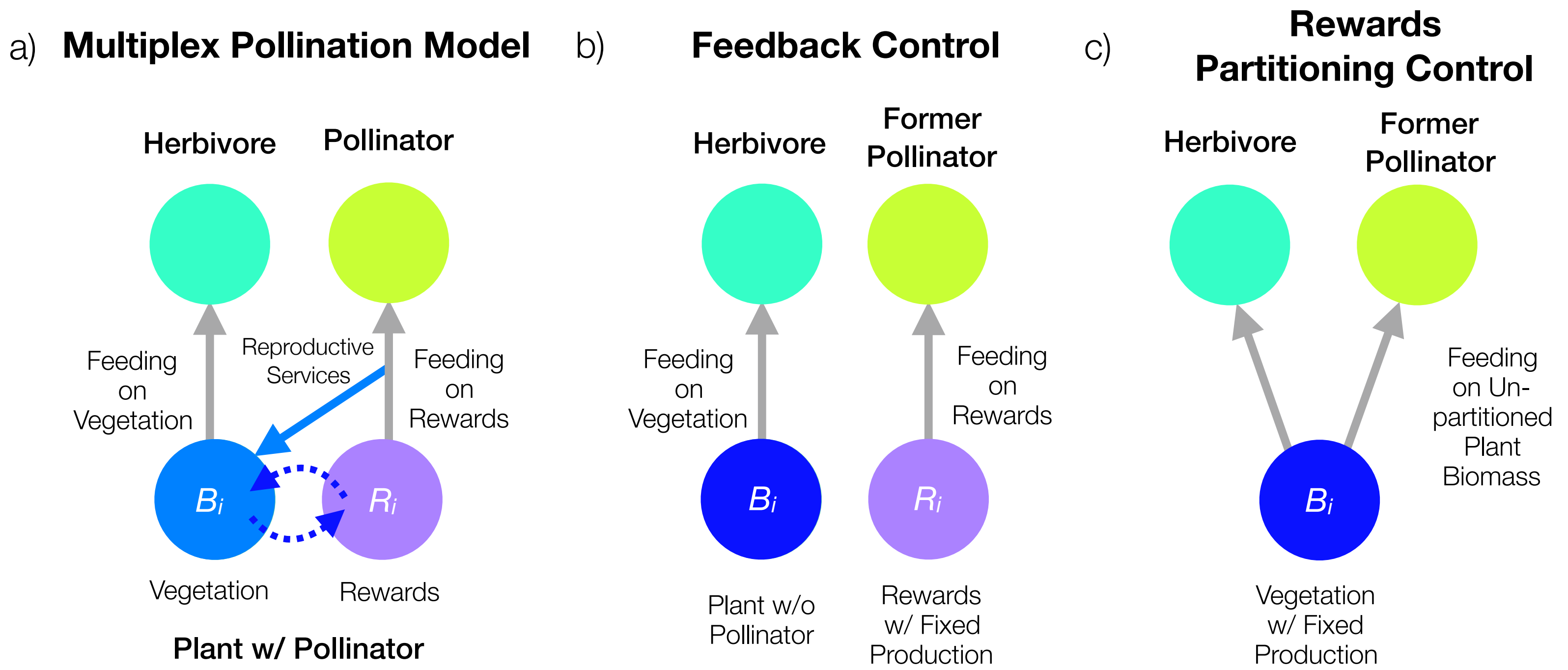

**Figure S7: Diversity, stability, and function in feedback controls.**

Full results for the feedback control treatments, in which food webs are initialized with rewards productivity from steady-state multiplex networks. Formatting of (a) and (b) follow main text Figs. 5 and 6, respectively, with light blue indicating *former* plants with pollinators. μ's corresponding to values reported in main text Fig. 4.

**(a)**

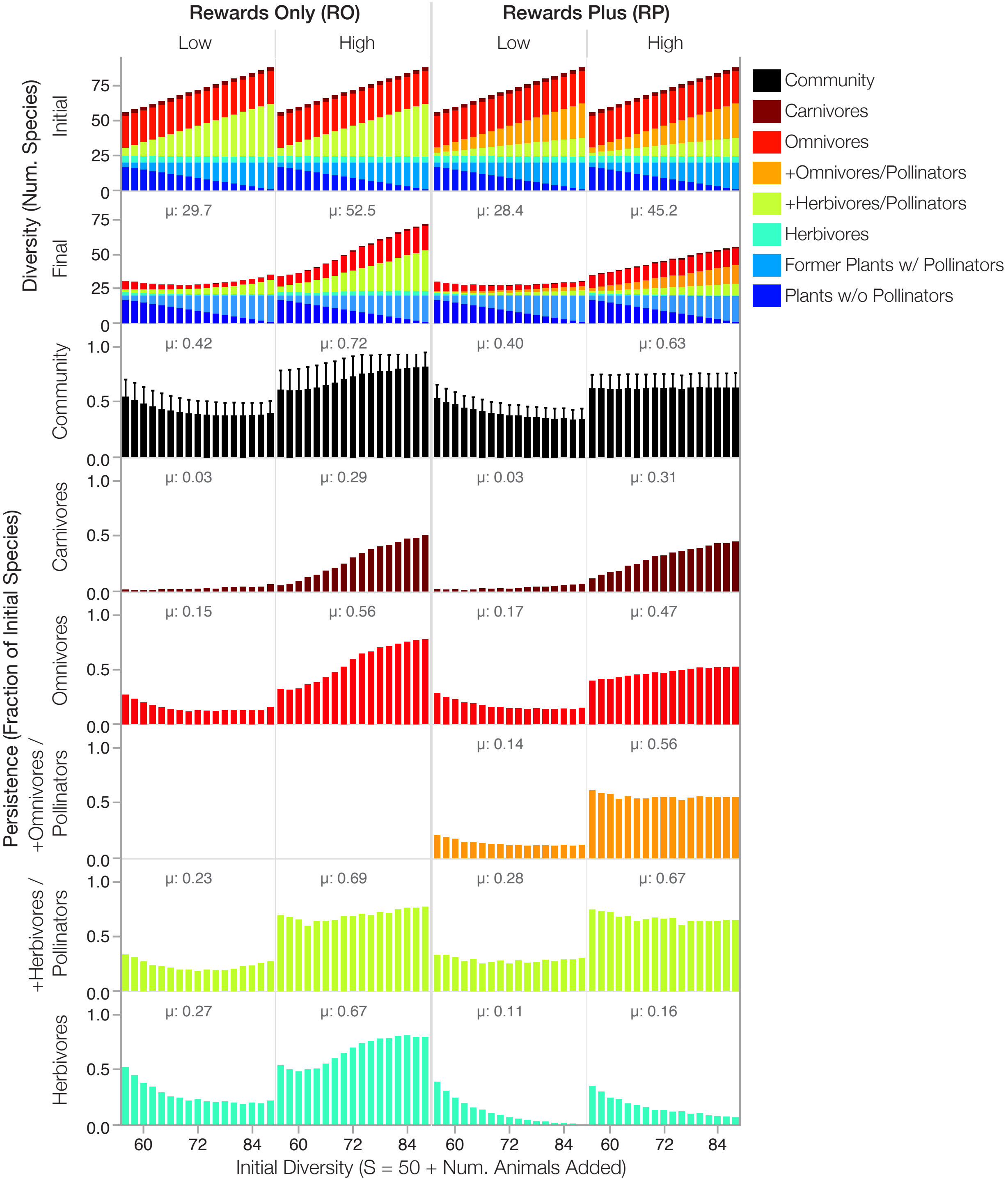

(b)

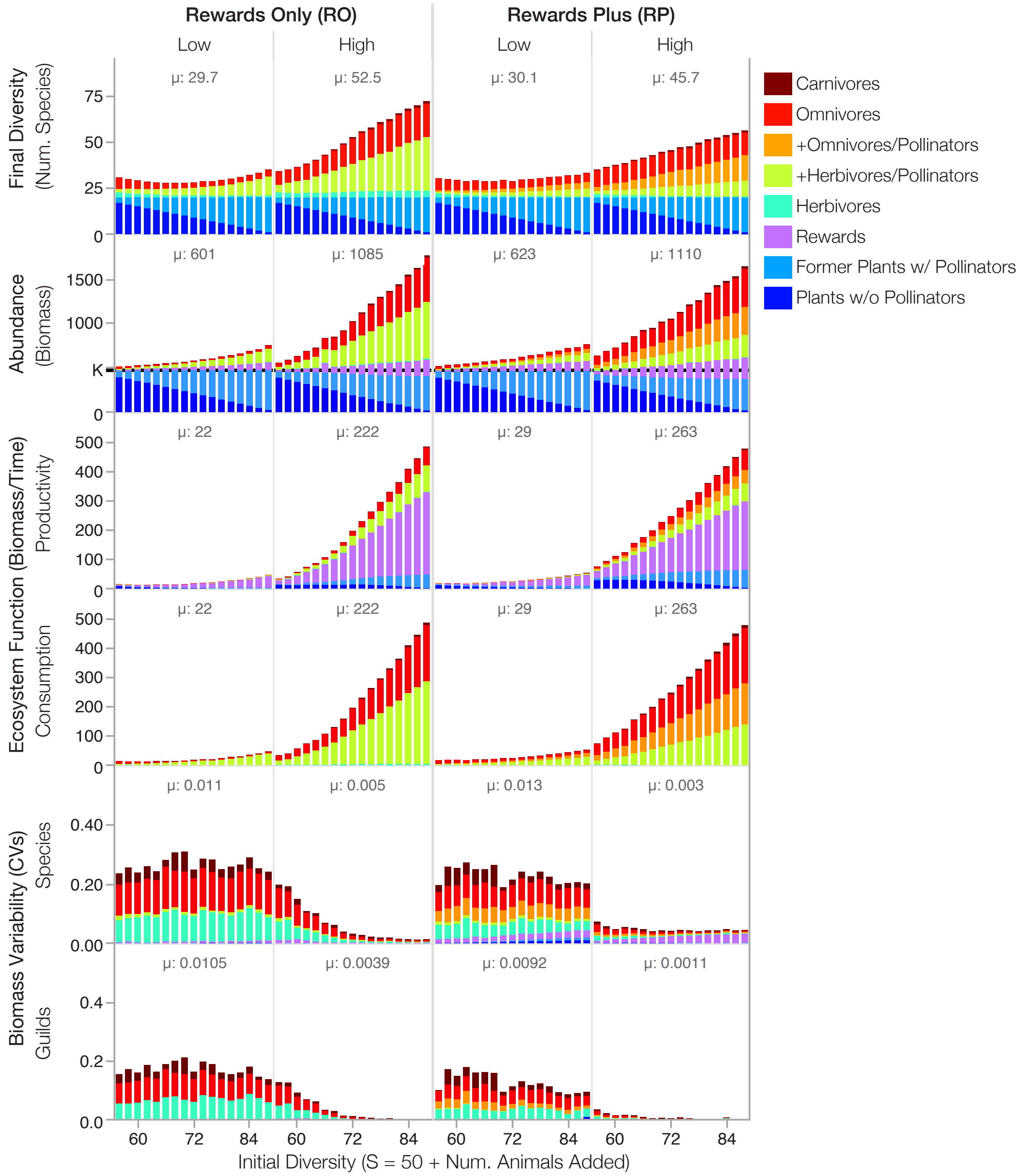
Rewards Only (RO)
Rewards Plus (RP)
Low
High
Low
High
Final Diversity (Num. Species)
μ: 29.7
μ: 52.5
μ: 30.1
μ: 45.7
Abundance (Biomass)
μ: 601
μ: 1085
μ: 623
μ: 1110
K
Ecosystem Function (Biomass/Time)
Productivity
μ: 22
μ: 222
μ: 29
μ: 263
Consumption
μ: 22
μ: 222
μ: 29
μ: 263
Biomass Variability (CVs)
Species
μ: 0.011
μ: 0.005
μ: 0.013
μ: 0.003
Guilds
μ: 0.0105
μ: 0.0039
μ: 0.0092
μ: 0.0011
Initial Diversity (S = 50 + Num. Animals Added)
Carnivores
Omnivores
+Omnivores/Pollinators
+Herbivores/Pollinators
Herbivores
Rewards
Former Plants w/ Pollinators
Plants w/o Pollinators

**Figure S8: Summary of community composition changes in feedback controls.**

Average absolute differences in steady-state a) persistence or c) total ecosystem biomass of the control simulations compared to their paired multiplex simulations. μ is the average over the $N$ = 24,276 simulations in each treatment. Though changes in ecosystem persistence and biomass are trivial, controls deviate systematically in the fraction of b) persisting species composed by each guild and d) total biomass composed by each guild. Positive values indicate positive effects of mutualistic feedbacks on persistence or biomass (corresponding to decreased persistence or biomass in controls compared to multiplex simulations). Error bars are 95% confidence intervals.

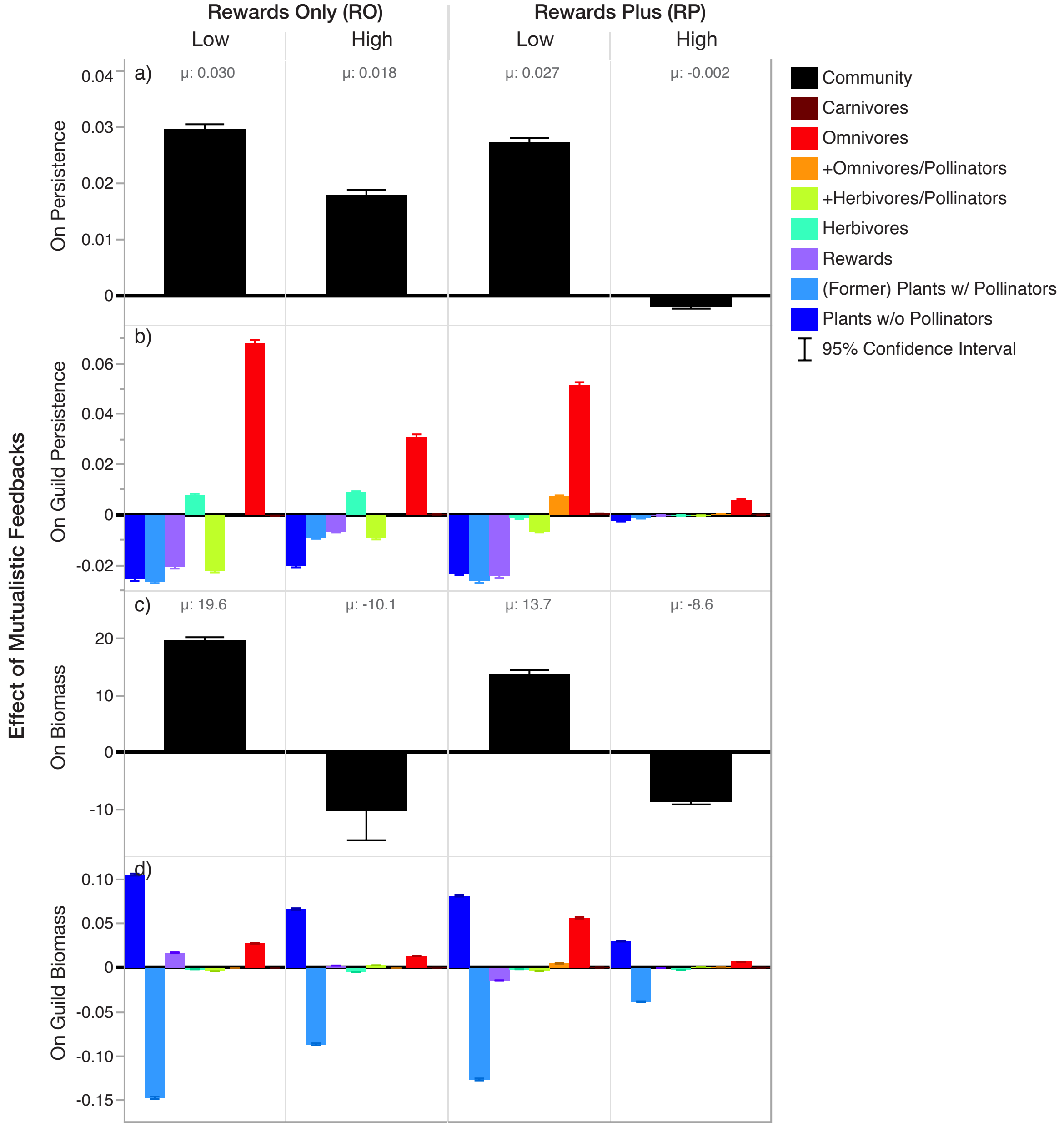

**Figure S9: Example timeseries comparing multiplex and feedback control simulations.**

This example uses a 50-species niche-model food web integrated with a 33-species plant-pollinator network following the same formatting as Fig. 3. Multiplex simulations (left columns) are compared to their paired feedback control simulations (right columns).

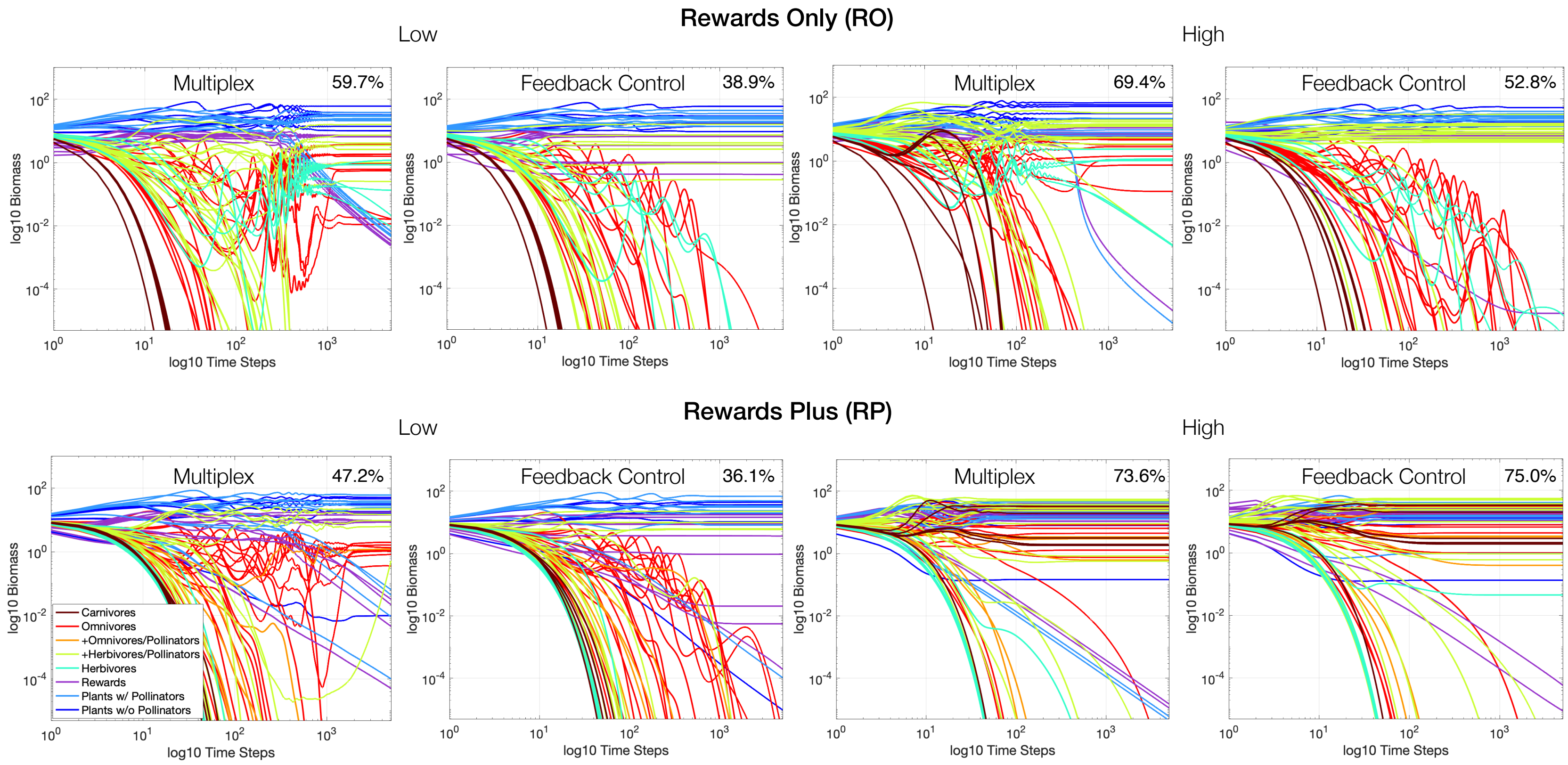

**Figure S10: Diversity, stability, and function in rewards partitioning control simulations.**

Full results for the rewards partitioning control treatments, in which steady-state multiplex networks are switched to dynamics of a traditional food web with unpartitioned rewards and vegetative biomass. Formatting of (a) and (b) follow main text Figs. 5 and 6, respectively, with light blue indicating *former* plants with pollinators. μ's corresponding to values reported in main text Fig. 4.

**(a)**

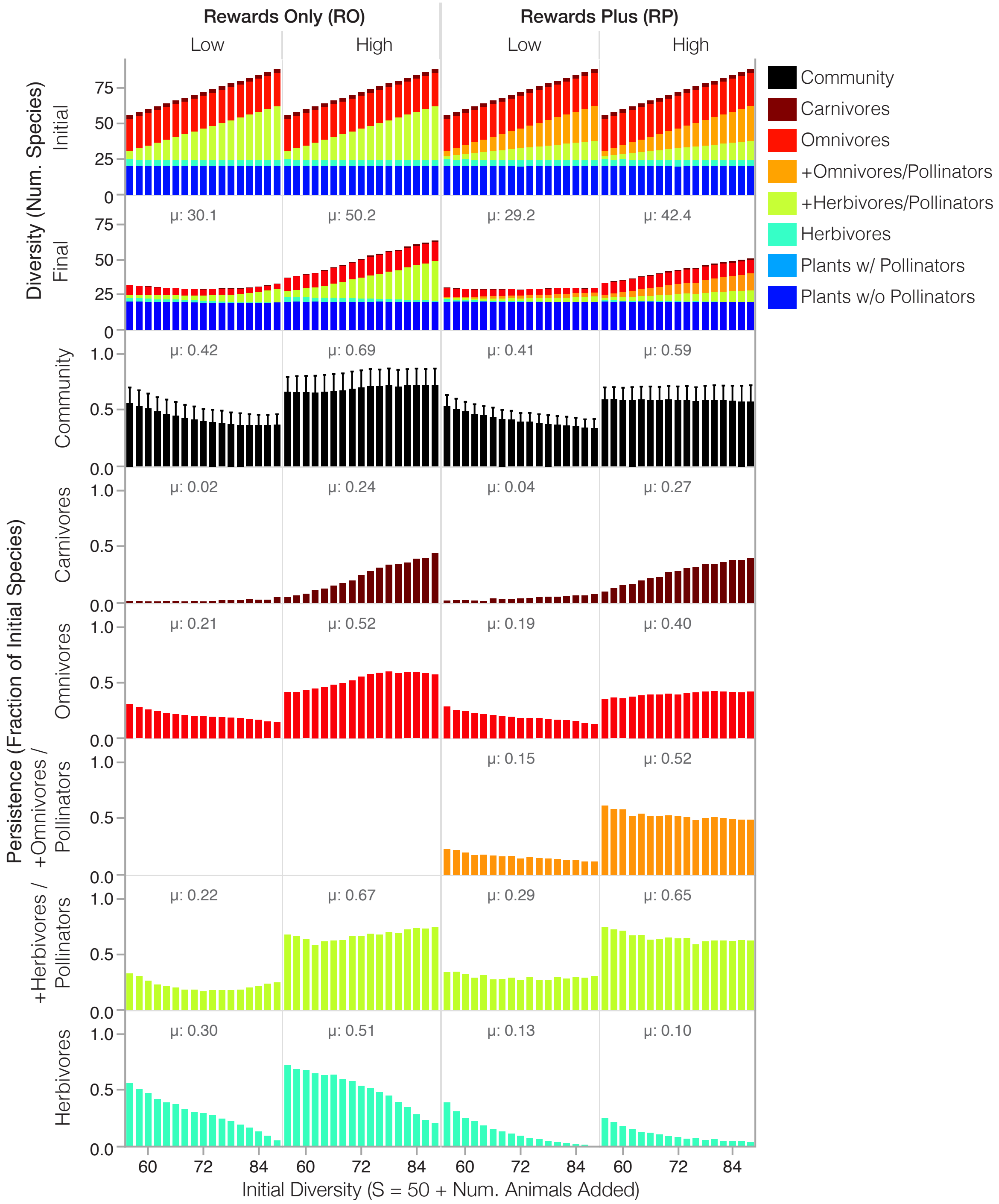

**(b)**

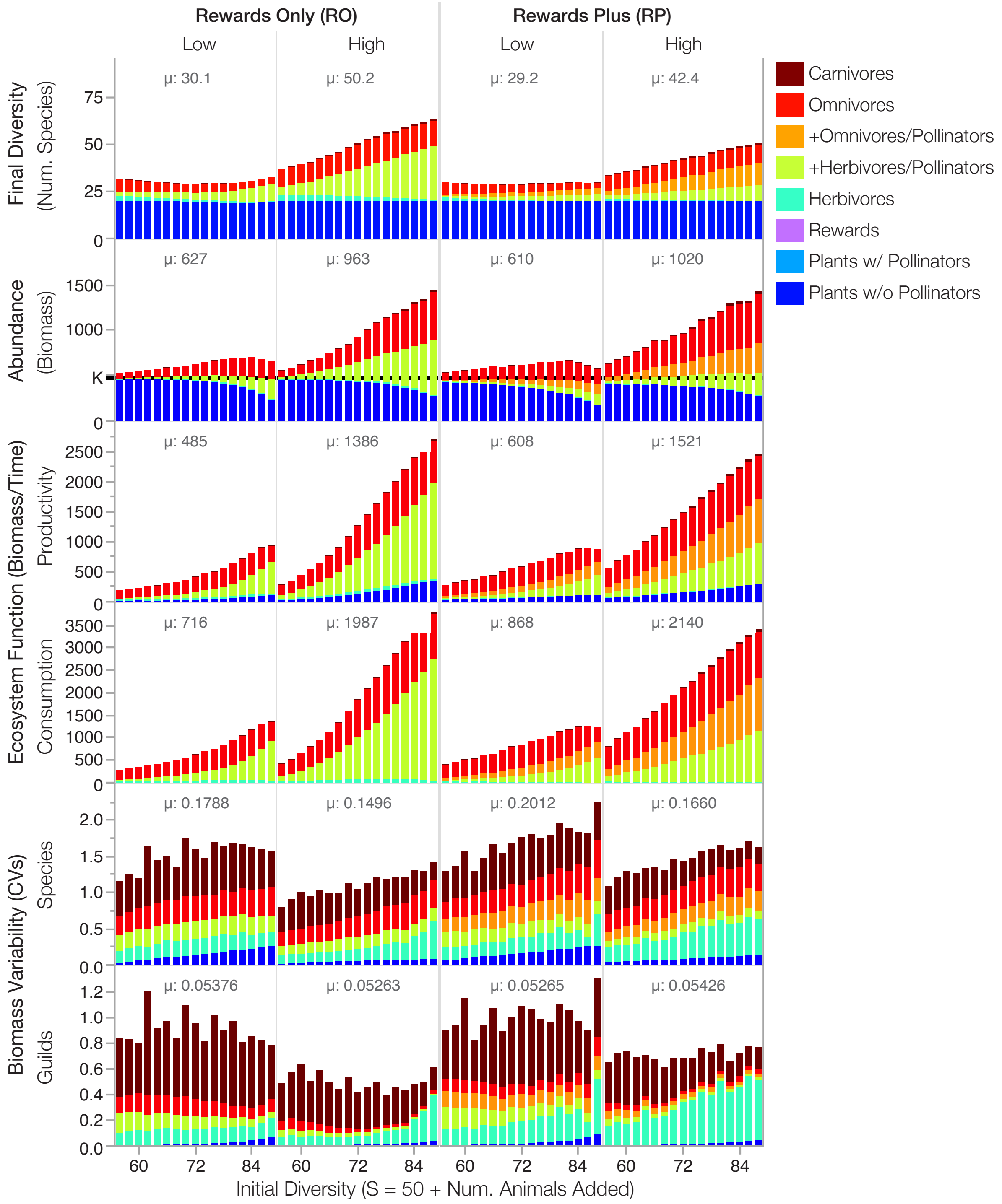
Rewards Only (RO)
Rewards Plus (RP)
Low
High
Low
High
Final Diversity
(Num. Species)
μ: 30.1
μ: 50.2
μ: 29.2
μ: 42.4
Abundance
(Biomass)
K
μ: 627
μ: 963
μ: 610
μ: 1020
Ecosystem Function (Biomass/Time)
Productivity
μ: 485
μ: 1386
μ: 608
μ: 1521
Consumption
μ: 716
μ: 1987
μ: 868
μ: 2140
Biomass Variability (CVs)
Species
μ: 0.1788
μ: 0.1496
μ: 0.2012
μ: 0.1660
Guilds
μ: 0.05376
μ: 0.05263
μ: 0.05265
μ: 0.05426
Initial Diversity (S = 50 + Num. Animals Added)
Carnivores
Omnivores
+Omnivores/Pollinators
+Herbivores/Pollinators
Herbivores
Rewards
Plants w/ Pollinators
Plants w/o Pollinators

**Figure S11: Summary of community composition changes in rewards partitioning controls.**

Results for the rewards partitioning control, following the formatting of Fig. S9.

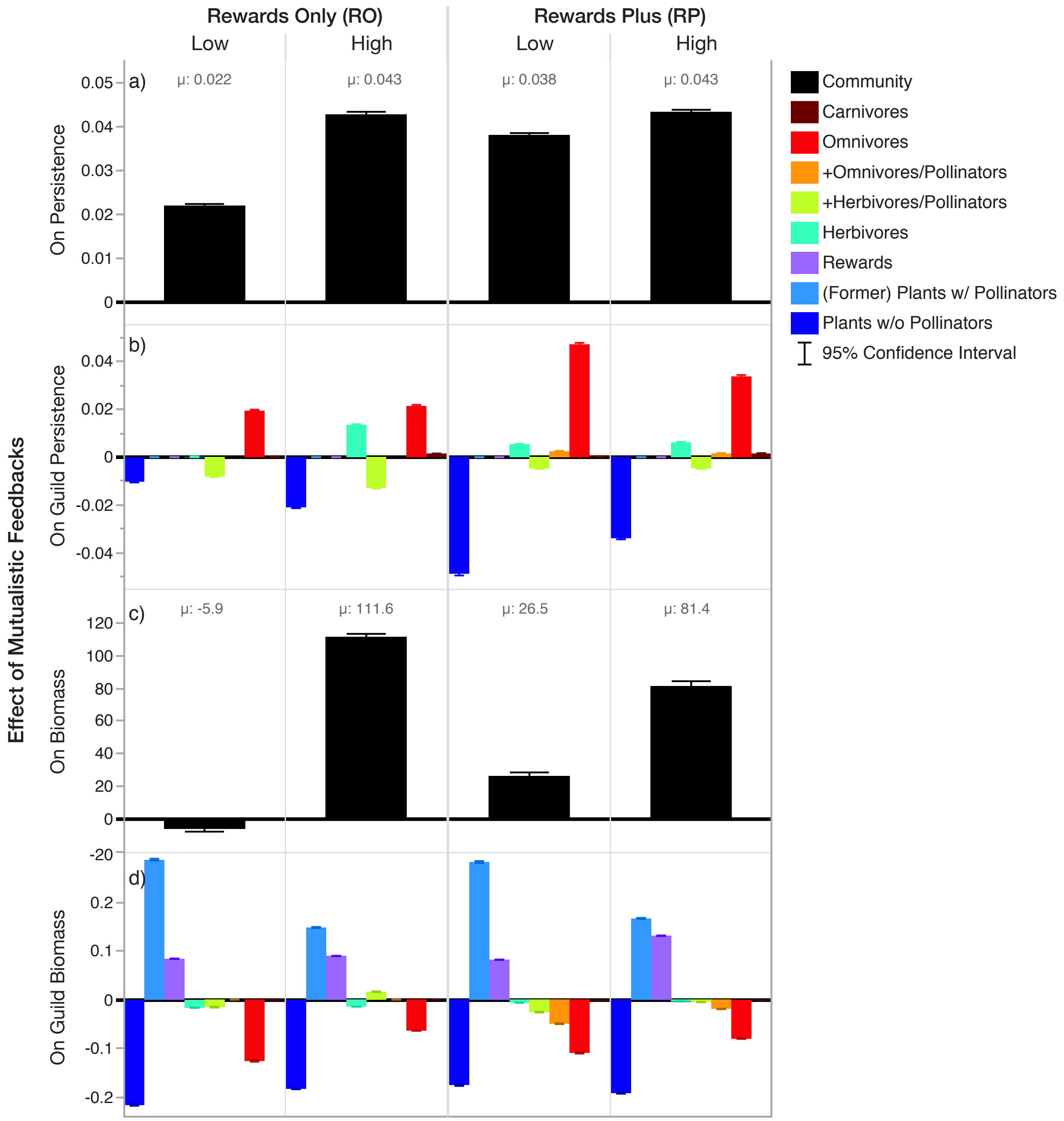
